# Conditional Generative Learning for Medical Image Imputation

**DOI:** 10.1101/2023.04.03.535422

**Authors:** Ragheb Raad, Deep Ray, Bino Varghese, Darryl Hwang, Inderbir Gill, Vinay Duddalwar, Assad A. Oberai

## Abstract

Image imputation refers to the task of generating a type of medical image given images of another type. This task becomes challenging when the difference between the available images, and the image to be imputed is large. In this manuscript, one such application, derived from the dynamic contrast enhanced computed tomography (CECT) imaging of the kidneys, is considered: given an incomplete sequence of three CECT images, we are required to the impute the missing image. This task is posed as one of probabilistic inference and a generative algorithm to generate samples of the imputed image, conditioned on the available images, is developed, trained, and tested. The output of this algorithm is the “best guess” of the imputed image, and a pixel-wise image of variance in the imputation. It is demonstrated that this best guess is more accurate than those generated by other, deterministic deep-learning based algorithms, including ones which utilize additional information and more complex loss terms. It is also shown the pixel-wise variance image, which quantifies the confidence in the reconstruction, can be used to determine whether the result of the imputation meets a specified accuracy threshold and is therefore appropriate for a downstream task.

## Introduction

Image imputation or image synthesis refers to the task of generating or synthesizing missing images using available data, which is often in the form of other types of images. In medical imaging, these techniques address tasks like generating images of one type (FLAIR MR image, for example) given images of a different type (T2 MR image), generating missing slices in a three-dimensional stack of slices of an organ, and generating an image at a specific time-point in a temporal sequence of images obtained from contrast-enhanced imaging modalities. Generally speaking, if the available images are “close” to the missing image, the image synthesis/imputation task is easier. For example, if the temporal sequence includes *≈* 50 images, wherein the sequential changes are small, then the missing image is easily approximated by a weighted sum of its neighbors in the sequence. On the other hand, if the available information is sparse, the image imputation task is challenging. For example, when the entire temporal sequence contains only a few images (say 4), and the difference between each image is significant, relying on neighboring images alone to infer a missing image is not an option. In this case, the image imputation algorithm must learn the complex dependencies between images in the sequence from an independent set of training data, and then apply this knowledge to the case of interest. These types of image imputation problems are the focus of this manuscript.

In particular, we consider the dynamic contrast enhanced computed tomography (CECT) imaging of the kidneys. In this modality, an intravenous contrast agent is injected into the subject and CT images are acquired at different time-points leading to pre-contrast, corticomedullary (30-40 seconds after injection), nephrographic (100 seconds after injection), and excretory (5-10 minutes after injection) phase images [20]. A complete sequence consists of all images at all four time-points. In some cases, one or more these images may be missing and may need to be imputed. For instance, subject motion during the exam could blur some images rendering them of little clinical value, or, under a clinical protocol a subject with a renal tumor might undergo CECT imaging where the pre-contrast image is not recorded. However, each image in the CECT sequence is important and has clinical value. For example, the pre-contrast, corticomedullary and nephrographic images are all required to evaluate intensity enhancement and washout within the tumor, kidney and other organs, which in is turn used in evaluation and diagnosis [6]. Also, in the excretory phase, the renal pelvis is clearly visualized and its location relative to the tumor can be determined. This information is useful to a surgeon performing nephrectomy to remove the tumor [3, 31].

Deep learning has found significant applications in tackling image synthesis/imputation problems of the type described above. Specifically, algorithms based on a U-net architecture that map images of one type to another via convolutional layers have been remarkably successful. Among these are a class of algorithms that typically employ adversarial learning and are closely related to generative adversarial networks (GANs) [10]. This includes the PIX2PIX algorithm which performs image transformation using pair-wise image data and adversarial loss augmented with a reconstruction term [15]. In CycleGAN [36] these ideas are extended by adding a cycle consistency loss and removing the stringent requirement of working with pairwise images. Algorithms like the StarGAN [7] and RadialGAN [35] extend these ideas to image transformation across multiple domains with a single generator network that uses a specific code which carries information about each domain. CollaGAN [18] provides a similar mapping across multiple domains by relying on multiple consistency losses. While these algorithms are based on probabilistic learning principles and have been remarkably successful in computer vision and medical imaging applications (see discussion below), they do not explicitly account for the the uncertainty in the image transformation task. Stated simply, given an image of one type they produce only one image of another type. They do not account for the fact that there may be an ensemble of images that are consistent with a given input image. Specifically, for the case of imputing CECT images, there may be whole collection of likely complete sequences that are consistent with a given incomplete sequence. The algorithm presented in this manuscript accomplishes this and doing so is shown to be more accurate and provide a measure confidence to its user.

In the probabilistic framework, we treat both the incomplete and complete image sequences as random vectors. Then, using data which consists complete sequences and their incomplete counterparts (generated by decimating an image at random), we train a conditional generative adversarial (cGAN) network [4, 22, 26]. This network takes samples from the joint distribution of two random vectors and learns to efficiently sample from the conditional distribution. That is, given an instance of one of these vectors, it generates samples of the other vector conditioned on that instance (see Figure 1). In our case, the “instance” is the incomplete CECT image sequence and the samples generated are the complete CECT sequences that are consistent with this incomplete sequence. From these samples we extract the desired imputed image, and compute the pixel-wise mean and standard deviation. The mean imputed image provides our best guess for the missing image, and the standard deviation image quantifies the uncertainty in our prediction. Through rigorous testing we show that the mean image is generally more accurate than images produced by methods that do not account for the probabilistic nature of the problem. We also demonstrate that the standard deviation image is correlated with the error in the imputation and can be used to quantify the confidence in the imputed image.

**Figure 1.**
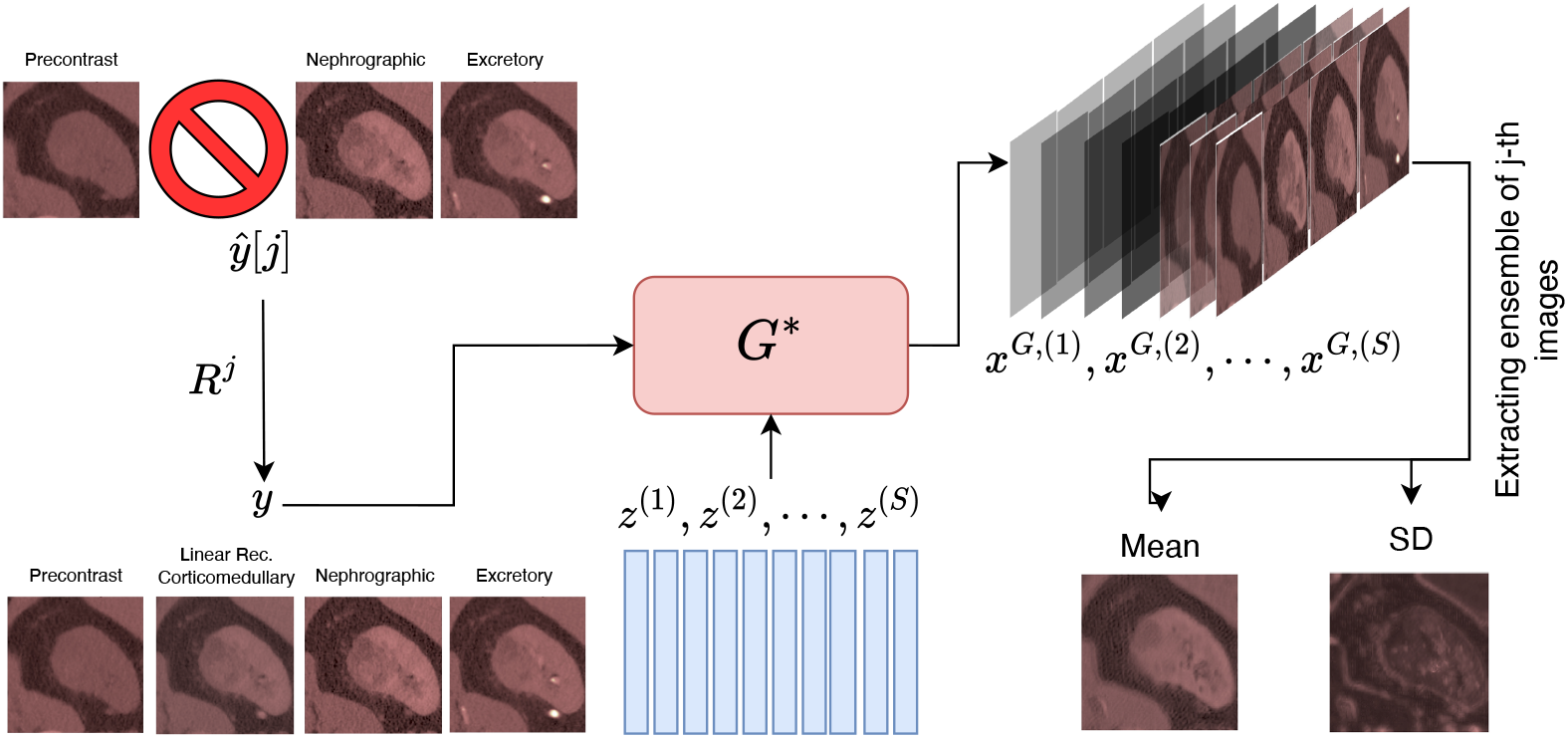
A schematic diagram of the imputation algorithm. In the above illustration, we have assumed that the corticomedullary phase is missing from the sequence (*j* = 2). The mean and standard deviation (SD) are evaluated using (2).

Deep learning-based image imputation techniques have recently been used for imputing and synthesizing CT images. This includes generating CT images for data augmentation to eventually improve the performance of a CT-image based classifier [9, 19, 30]. It also includes algorithms for generating CECT images at a single time point for the lungs [21], and the kidneys [27], where the latter study uses the concept of neural transfer for improved performance. Recently, several GAN-based approaches were implemented and tested for imputing renal CECT images at different time-points [32]. The approaches tested include several standard algorithms and two novel methods, ReMIC and DiagnosisGAN, which were shown to be the most accurate. In ReMIC [28], a representational disentanglement scheme for multi-domain image completion was used to improve the performance of the algorithm, whereas in DiagnosisGAN [32] in addition to the CECT images themselves, other sources of information, like segmentation mask for the tumor, and the knowledge cancer sub-type, were used to improve the performance of the method. In the results section of this manuscript we compare our algorithm with these methods and conclude that our method is more accurate, and at the same time provides estimates of confidence in the imputation task.

We remark that in an earlier work [25] we presented a probabilistic method for imputing CECT images where we utilized GANs to learn the prior distribution [23, 24] of a complete sequence of CECT images. This prior was combined with a likelihood term driven by a measured incomplete sequence to set up a Bayesian estimate for the probability distribution of the complete sequence. This inference problem was then solved by advanced Markov-Chain Monte Carlo (MCMC) methods. In the present approach, in contrast to this, we utilize a conditional GAN to directly learn and sample from the conditional distribution. This leads to an an algorithm that is much more efficient. As a result, we have applied it to reconstructing slices with the tumor, the kidney and the surrounding tissue, whereas the earlier work was limited to segmented images of just the tumor.

The remainder of this manuscript is organized as follows. In the following section, we present the results obtained by applying the our algorithm to incomplete sequence of CECT images. Thereafter, we discuss these results and their medical relevance. Finally, in the Methods Section we describe our algorithm in detail.

## Materials and Methods

We begin this section by introducing some mathematical notations necessary to explain how a cGAN is used to solve the imputation problem in a Bayesian framework. Define the product space 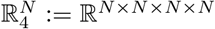, where *N* = 128 *×* 128 is the size/resolution of each image. 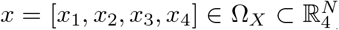 be the sequence of CECT images for a patient at the four time-points. Let 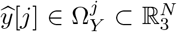 be the patient’s CECT image sequence with the *j*-th image in the sequence missing. The missing image is replaced by a simple linear reconstruction using map 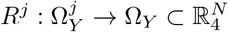 with a mapping existing for each 1*≤ j≤* 4 (see Supplementary Note 1). Note that the linear reconstruction of the missing image only serves as an initial guess, and is typically unable to represent several desired features and intensity variations of the true image. We use the notation 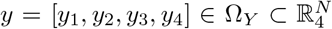 to denote the final measurement where an incomplete sequence has been filled in with the linear approximator. We are interested in finding an appropriate *x* given the measurement *y*.

To accommodate for the fact the reconstructed *x* may not be unique, we consider a Bayesian formulation where sequences *x* and *y* are modelled using random variable *X* and *Y*, respectively. We are thus interested in finding the conditional probability distribution *P*_*X*|*Y*_ given an incomplete measurement *Y* = *y*, and generating samples from this distribution. Furthermore, we want to learn this distribution by only working with a finite set of paired samples *{x*^(*i*)^, *y*^(*i*)^ *}* drawn from the joint probability distribution *P*_*XY*_. This is achieved using a cGAN which comprises two networks, namely the generator *G* and the critic *D*. The generator is given by the mapping *G* : Ω_*Z*_ *×*Ω_*Y*_ *→*Ω_*X*_ with *x*^*G*^ = *G*(*z, y*), where *z* is a realization of the *N*_*Z*_-dimensional latent random variable *Z* defined on Ω_*Z*_. The latent vector is typically chosen to follow a simple distribution *P*_*Z*_, such as a multivariate Gaussian, which is easy to sample from. The role of the *G* is to generate samples (given *Y* = *y*) from the learned distribution 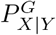 which are similar to samples from the true conditional. The critic is given by the mapping *D* : Ω_*X*_*×* Ω_*Y*_*→* ℝ, with its role being to distinguish between true joint samples (*x, y*) *∼P*_*XY*_ and fake joint samples (*x*^*G*^, *y*) where *x*^*G*^ is a fake sample generated by *G*.

We consider a particular cGAN variant, known as the Wasserstein cGAN [4], which makes use of the following loss function

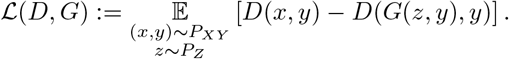

The two networks are trained simultaneously by solving the following minmax problem

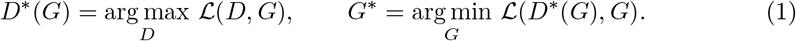

Under the assumption that the critic is 1-Lipschitz and that the maximization problem is solved perfectly, it can be shown that finding the optimal generator is equivalent to minimizing the mean (with respect to the marginal distribution *P*_*Y*_) Wasserstein-1 distance between *P*_*X*|*Y*_ and 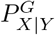 [4]

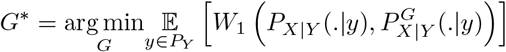

where *W*_1_ is Wasserstein-1 metric [33]. The Lipschitz constraint on the critic can be weakly imposed using a gradient penalty term while training *D* [4, 26] (also see Supplementary Note 3).

*In practice, we do not know the exact form of P*_*XY*_, however, we assume access to samples 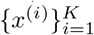 of complete image sequences drawn from the marginal *P*_*X*_ of *X*.

For each of these samples, we drop the *j*-th image in the sequence and use the linear reconstructor *R*^*j*^ to construct the corresponding *y*. Note that four such *y*’s can be constructed for each *x*. This leads to the dataset 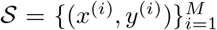 with *M* = 4*K* samples, where each pair can be seen as a sample from the joint distribution *P*_*XY*_. Using these samples, the expectations in () are approximated by empirical averages.

Once the cGAN is trained, given a new incomplete measurement 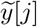 we can use the trained *G*^*∗*^ to generate an ensemble of probable *x*’s and evaluate their pixel-wise statistics

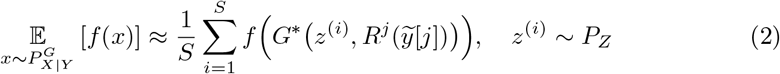

for any continuous, bounded function *f* on Ω_*X*_. By setting *f* (*x*) = *x* in (2) we can evaluate the mean prediction denoted by 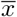, which serves as our best guess for the complete sequence. Choosing 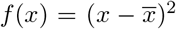in (2), we can evaluate the pixel-wise variance of the learned posterior distribution. This variance (or rather the standard deviation) can be used to quantify the uncertainty in the reconstructed sequence. The schematic of the reconstruction algorithm is shown in Figure 1. Note that if the *j*-th image is missing, we are typically only interested in statistics of the *j*-th images of the generated ensemble.

### cGAN architecture

The architecture of the generator and critic is based on those considered in [26]. The generator *G* has a U-Net architecture, as shown in Figure 2(a), taking as input the measured sequence *y* (after linear reconstruction of the missing phase) and the latent variable *z*. The latent information is injected at every scale of the contracting and expanding branches of the U-Net using conditional instance normalization (CIN) [8], which has two advantages: i) the latent dimension *N*_*Z*_ can be chosen independently of *N*_*X*_ or *N*_*Y*_, thus allowing for significant dimension reduction, and ii) stochasticity is introduced at all scales of the U-Net, which overcomes the issue of mode collapse [26].

**Figure 2.**
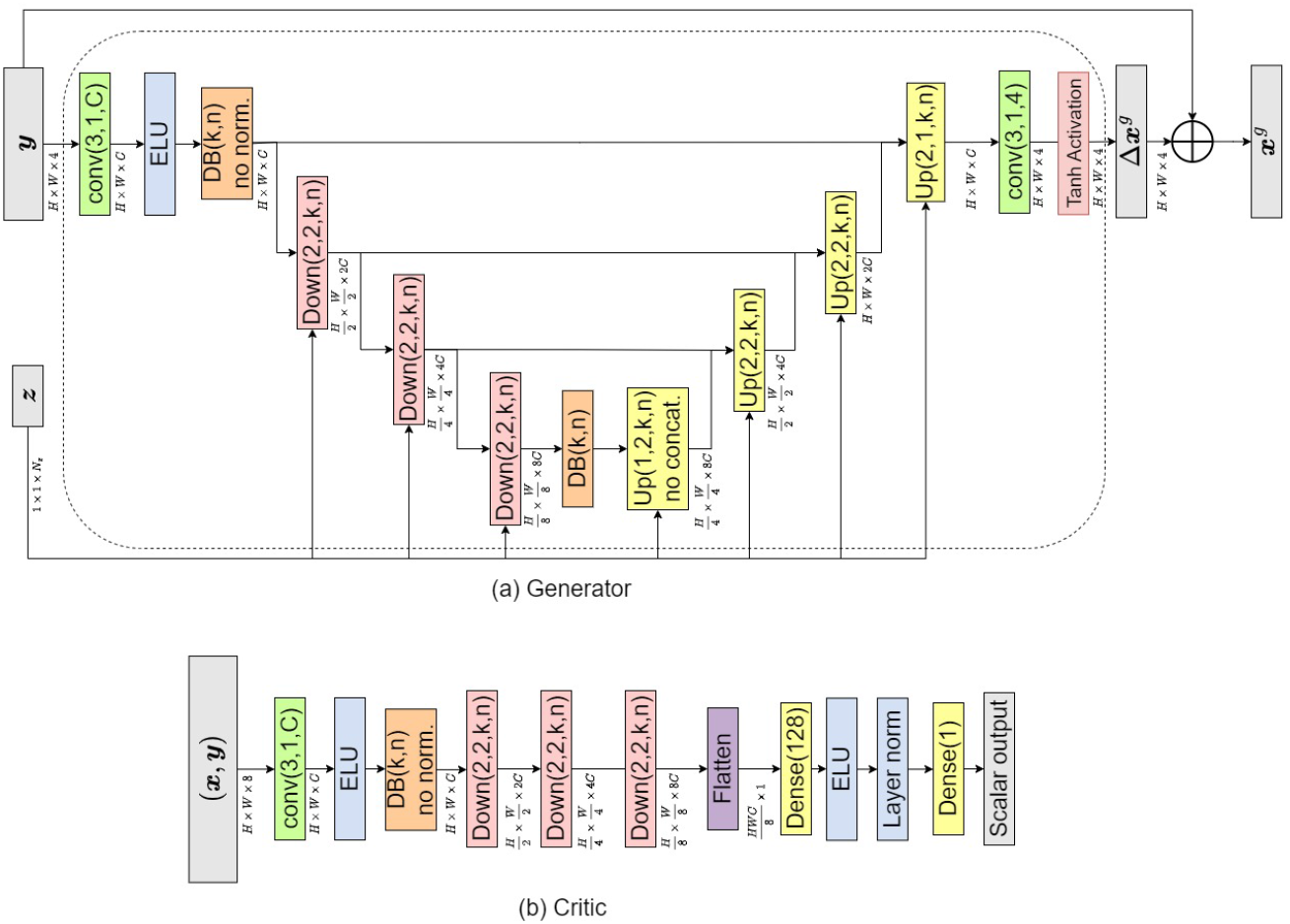
Architecture of generator and critic used in the conditional GAN.

It is typical to use residual blocks to introduce non-linearity in the U-Net, which is what was also done in [26]. However, in the present work, we make use of dense blocks since they lead to superior performance compared to residual blocks while reducing the number of trainable parameters [13]. In Figure 2, the dense blocks are denoted by DB(*k, n*), where *n* corresponds to the number of sub-blocks in the dense block, while *k* denotes the number of output features in all but the last sub-block. Down(*p, q, k, n*) denotes a down-sampling block, which coarsens the input spatial resolution by a factor of *p*, while increasing the number of channels by a factor of *q*. The parameters *k, n* correspond to the dense block used in the down-sampling block. Similarly, Up(*p, q, k, n*) denotes the up-sampling block, which refines the spatial resolution by a factor of *p* and decreases the number of channels by a factor of *q*. Further details about the constitutive blocks of the U-Net are given in the Supplemental Note 2.

Another difference between the present architecture as compared to the U-Net in [26] is the use of an outer skip connection, which adds the output of the U-Net to the measurement sequence *y*. Thus, the U-Net’s output can be interpreted as a pixelwise perturbation ∆*x*^*G*^ of all the images in the sequence, which is added to the input measurement *y* to predict a complete sequence *x*^*G*^. Such skip connections are routinely used in U-Nets trained for image de-noising applications [11, 12].

As shown in Figure 2(b), the architecture of the Critic *D* comprises dense block-based down-sampling, followed by fully connected network that gives a scalar output. Since the latent variable is not used to evaluate the critic, CIN is replaced by layer normalization [5]. The original Wasserstein cGAN [4] use a more complicated and specialized critic to overcome the mode collapse. However, as discussed and demonstrated in [26], the injection of sufficient stochasticity in the generator via CIN allows the use of a simpler critic architecture.

### PIX2PIX: A non-stochastic algorithm

To demonstrate the benefits of a stochastic imputation model, we compare the results of the proposed cGAN with a PIX2PIX GAN based on the work described in [15]. This is considered the standard model to use in image-to-image translation. The PIX2PIX model in our paper is modified to resemble our cGAN model for a fair comparison. The generator of the PIX2PIX approach is required to not only fool the critic but to also make a prediction that is (point-wise) close to the ground truth. This motivated the augmentation of *l*_1_ distance term in the generator loss [15], which is the approach we follow as well. The generator for the PIX2PIX model is given by the map *G* : Ω_*Y*_ *→*Ω_*X*_ and does not use a latent variable. Thus, a single *x*^*G*^ is generated for a given *y*, unlike our cGAN model. The loss function for PIX2PIX is given by

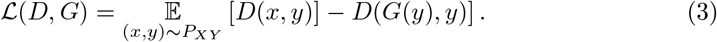

To train PIX2PIX, the following minmax problem is solved

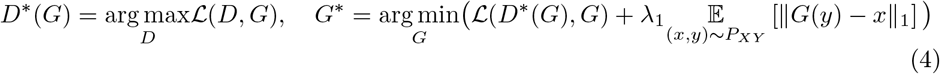

The architectures of the networks will be similar those used in the cGAN approach with the exception that batch normalization [14] is used instead of CIN in the generator.

### Other generative models

We briefly describe two existing deep generative models that have been developed for medical image imputation tasks. The first is DiagnosisGAN [32], which is a specialized GAN model that can simultaneously generate missing image in CT sequences and classify the cancer subtype. In this model, the generator take an incomplete multi-phase CT sequence as input and generates a synthesized volume for the missing phase. The training objective function is composed of several loss terms, including the adversarial loss, a reconstruction loss, a lesion segmentation loss, and cancer subtype classification loss.

Another approach is known as the Representational disentanglement scheme for Multi-domain Image Completion (ReMIC [28]), which is a multi-domain completion and segmentation framework. The ReMIC model consists of a content encoder, which is shared across all domains (FLAIR or MRI for example). There are also domain-specific style encoders, and generators. In addition to an adversarial loss, the framework includes an image consistency loss for visible domains, latent consistency loss, and reconstruction loss for the generated images. Additionally, the framework employs a representational learning approach, where a segmentation generator follows the content code for a unified image generation and segmentation.

We remark that compared to these two models, the proposed cGAN has a much simpler architecture and objective function. Further, the cGAN generator is capable of generating an ensemble of possible missing images for an incomplete CT sequence instead of a single reconstructed image. The ensemble statistics enable us to quantify the uncertainty in the reconstruction.

## Results

### Patient Data

The study population consists of patients who had renal masses diagnosed on abdominal CECT scans and underwent resection at USC between May 2007 and September 2018. The pathology of the masses was confirmed after resection, and the patients were identified through a query of a surgical database. Patients without evaluable preoperative imaging or missing any of the four time-points of the CECT study were excluded. The final cohort included 370 patients, and three-dimensional regions of interest of the renal masses were manually segmented by two senior radiologists using Synapse 3D software [2]. The original images were 512 *×* 512 pixels with a pixel size of 0.9765 mm in each direction. These images were cropped to a size of 128*×* 128 by selecting a square centered on the tumor centroid. From the total data, 296 subjects were used for training (around 80%), 37 subjects were used for validation (around 10%), and 37 subjects were used for testing the algorithm (around 10%). The training data was augmented by rotating each image by *±*10, *±*20 and *±*30 degrees and by shifting it in the horizontal and vertical directions. This yielded at most eleven images for each original image. Data from Hounsfield units (within the range (*−*3024, 3071)) was normalized to the range (*−*1, 1) by first clipping values below -150 and above 1000 and then linearly transforming the Hounsfield scale to the normalized range.

### Image Imputation Results

Figures 3-5 display the results of the image imputation algorithm for six subjects selected from among the 37 test subjects. These subjects were selected to highlight the diversity of the type of renal CECT images that the algorithm can be applied to. Results for all 37 subjects are available at [1]. Each figure represents results from two subjects, and for each subject, the first row displays true images, the second row contains the mean imputed images generated by the cGAN, and the third row contains the images of the pixel-wise standard deviation generated by the cGAN. The columns represent the four time-points of the CECT exam. The mean imputed image may be interpreted as the “best guesss” generated by the cGAN, while the standard deviation image represents the spatial distribution in the uncertainty in this imputation.

**Figure 3.**
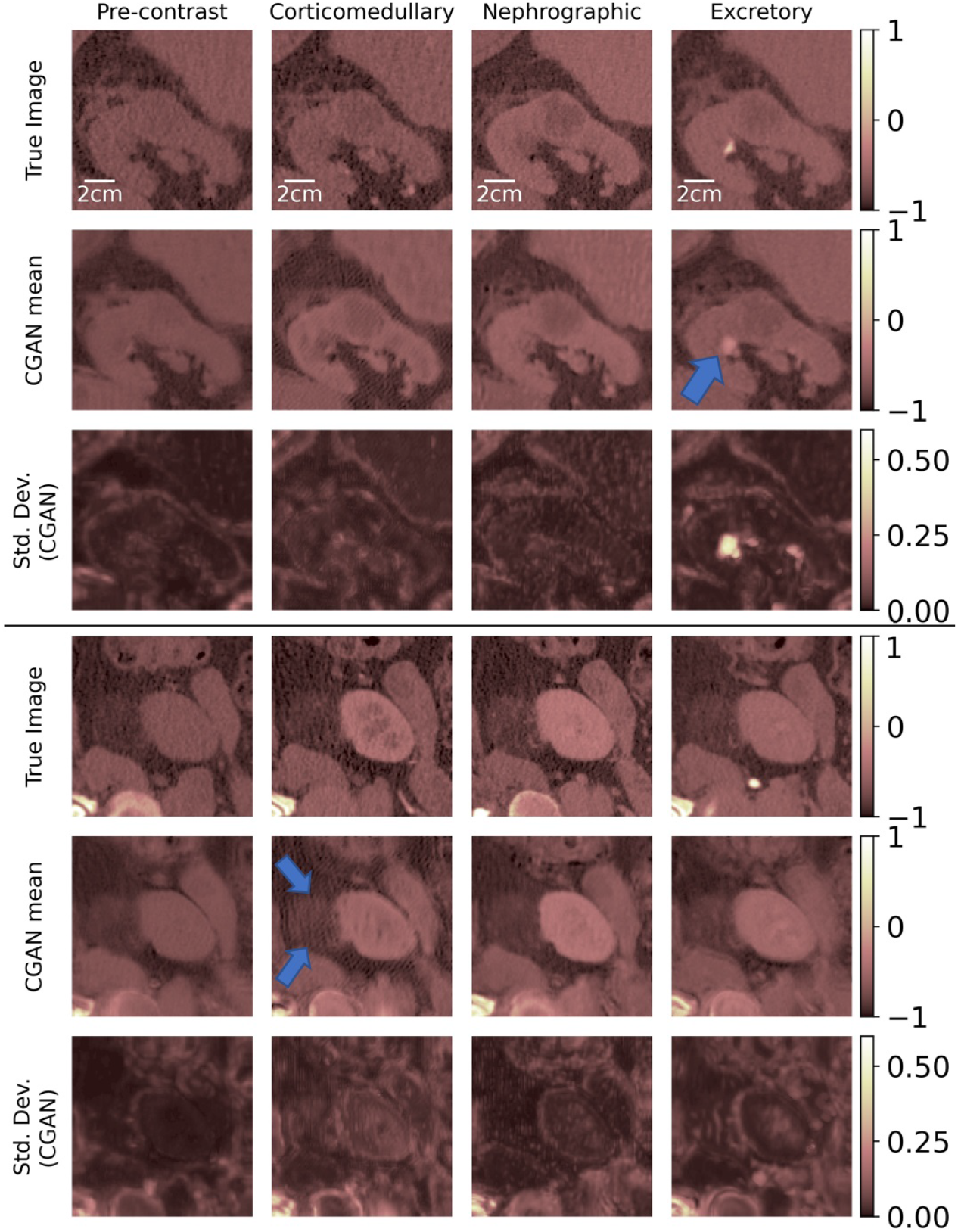
True and imputed images for Subjects 1 and 2.

To generate the imputed images, the algorithm assumes that the true image for a specific time-point is missing and needs to be predicted by making use of the images at the three other time-points. For example, for a given subject, images in the first column in rows 2 and 3 are generated by assuming that the true image for the pre-contrast time-point is missing and needs to be imputed by using the true images at the other three time-points. Similarly, images in the second, third, and fourth columns are obtained by imputing missing images for the corticomedullary, nephrographic, and excretory time-points, respectively, while making use of the true images from complementary set of time-points. Therefore, each figure demonstrates the capacity of the cGAN algorithm to impute images for the four distinct CECT time-points.

In Table 1, we perform a quantitative comparison of the performance on the cGAN with an improved variant of the popular PIX2PIX [15] algorithm (described in the Methods Section) which may be thought of as a deterministic version of the cGAN approach. We implement this method, and train and test it using the same image data used for the cGAN. From this Table it is clear that the cGAN method outperforms the PIX2PIX algorithm for all time-points. It is also interesting to note that it is most accurate in imputing pre-contrast images and least accurate in imputing corticomedullary images.

**Table 1.** Performance of the cGAN and PIX2PIX algorithms on test data (values averaged over 37 subjects).

|  |  |  |  |  |
| --- | --- | --- | --- | --- |
| SSIM | Pre-Contrast | Corticomedullary | nephrographic | Excretory |
| cGAN | **0.7262** | **0.6232** | **0.6353** | **0.6607** |
| PIX2PIX | 0.7022 | 0.6057 | 0.5992 | 0.6111 |
| PSNR | Pre-Contrast | Corticomedullary | nephrographic | Excretory |
| cGAN | **28.43** | **26.66** | **26.95** | **25.16** |
| PIX2PIX | 27.25 | 25.84 | 26.02 | 23.89 |
| NMSE | Pre-Contrast | Corticomedullary | nephrographic | Excretory |
| cGAN | **0.1043** | **0.1302** | **0.1297** | **0.1587** |
| PIX2PIX | 0.1184 | 0.1472 | 0.1457 | 0.1832 |

In Table 2, we compare the performance on the cGAN and PIX2PIX algorithms implemented in our study with the performance of the two top-performing algorithms reported in a recent CECT renal image imputation study [32]. These algorithms are

**Table 2.** Performance of different algorithms averaged over all time-points and subjects. Each entry contains the average and the standard deviation (in parenthesis).

|  |  |  |
| --- | --- | --- |
| Model | SSIM | PSNR |
| cGAN | **0.6614 (0.0845)** | **26.80 (2.91)** |
| PIX2PIX | 0.6295 (0.0927) | 25.75 (3.05) |
| DiagnosisGAN [32] | 0.6243 (0.1045) | 20.07 (2.06) |
| ReMIC [32] | 0.6333 (0.1012) | 20.33 (2.02) |

1. DiagnosisGAN [32], which is a generative adversarial algorithm like the cGAN. However unlike the cGAN approach, it does not provide any information regarding the confidence in the imputation and further it requires segmented tumor images to improve its performance.

2. ReMIC [28], which relies on a representational disentanglement scheme for multi-domain image completion. When compared with the cGAN approach, the ReMIC algorithm is more complex and relies on different types of losses which include image and latent domain consistency losses, adversarial losses, and reconstruction losses. In contrast to the cGAN approach, the ReMIC approach is also a deterministic approach and does not yield any information regarding confidence in the imputed images.

The interested reader is referred to the Methods section and the original references for further details on these methods. From Table 2, which reports the values of the normalized mean squared error (NMSE), the structural similarity index measure (SSIM) and the peak signal-to-noise ratio (PSNR) averaged over all time-points and all subjects, we conclude that for both these metrics the cGAN approach is the most accurate.

### Standard Deviation and Uncertainty

While the cGAN-based method provides more accurate imputation results than other methods, its true benefit is its ability to draw an ensemble from the posterior distribution rather than just a single most likely sample. This ensemble can be utilized to compute statistics that offer insight into the confidence in the imputed image. We demonstrate this by computing the estimated pixel-wise standard deviation in the imputed images in Figures 3-5 (last row). These images offer a spatial depiction of the level of uncertainty present in the imputed results. The higher the value of standard deviation in zone of pixels, the larger the ensemble variation and uncertainty in that zone.

We investigate the relation between the standard deviation computed by our algorithm and the true error of the image imputed by the cGAN. If a positive correlation is identified, then the standard deviation may be utilized as an indicator of the error in the imputed image, thus serving as a powerful tool for the end-user to eliminate imputed images that are likely to be inaccurate. In particular, we determine whether the total value of standard deviation (summed over all pixels) can be used to classify a given imputed image as being acceptable. We set a threshold of NMSE = 0.1 as a criterion and bin each imputed image into “acceptable” and “not acceptable” classes. Thereafter, we quantify the performance of the total standard deviation as a surrogate for performing this classification. The results are summarized in the receiver operating characteristic (ROC) curve shown in Figure 6. For this curve, the corresponding the area under the curve (AUC) value is 0.8825. Based on this we conclude then that the total standard deviation is predictive of whether a given imputation is sufficiently accurate.

## Discussion

In images for Subject 1 (Figure 3) we observe a well marginated tumor which is predominantly endophytic (growing within the kidney rather than protruding out) and has density similar to that of soft tissue. This tumor was a surgically proven renal cell carcinoma. The generated images demonstrate the lesion and the margins at each time-point. The subtle nodular peripheral enhancement noted in the true corticomedullary and nephrographic phases is also clearly seen in the generated images. In addition, in the imputed excretory phase image the location of the adjacent calyx (arrow) where urine collects from that portion of the kidney is anatomically consistent with the true image. We note that the depiction of the tumor, its margins and its relationship with the calyx and other intra-renal structures are important for surgical planning. All these features are well reproduced in the imputed images.

In images for Subject 2 (Figure 3), we observe a patient with an exophytic (protruding out of the kidney) tumor which was an angiomyolipoma. The imputed images reproduce the density of the tumor and its relationship with the kidney very well. The depiction of the density of the tumor, viz. a density that is predominantly the same as the density of fat tissue, enables the accurate diagnosis of this specific tumor via imaging.

The tumor in Subject 3 is a complex cystic (with roughly the density of a fluid) mass with multiple irregular nodular enhancing septations within it (Figure 4). The complexity, nodularity of the septations are important diagnostic features used by radiologists to characterize cystic tumors using the Bosniak scoring system [16, 17, 29, 34]. This was a Bosniak 4 tumor indicating (and proven) malignant cystic renal carcinoma. We note that these important features are reproduced in the imputed images.

**Figure 4.**
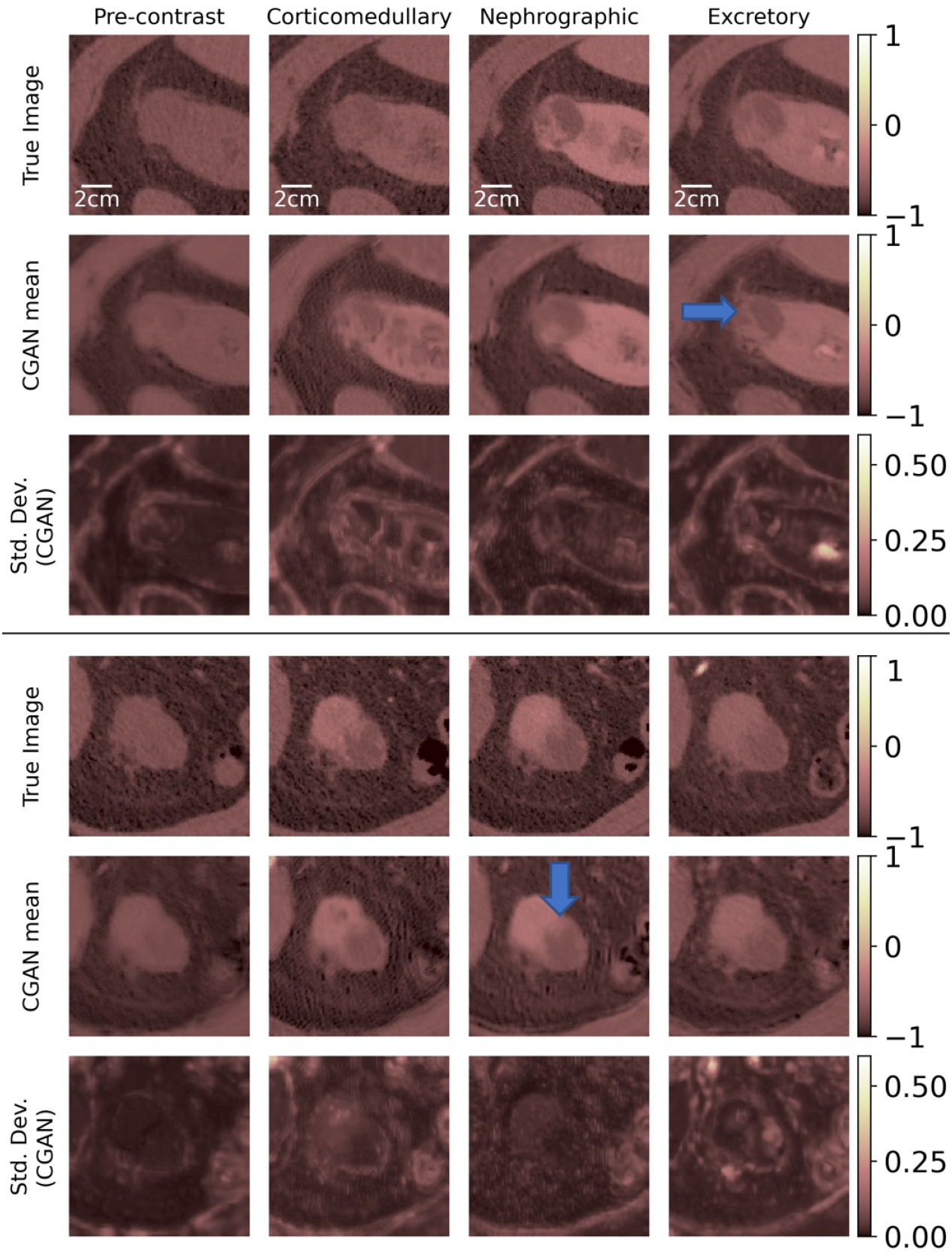
True and imputed images for Subjects 3 and 4.

Subject 4 is another example of a complex cystic tumor where the nodular peripheral enhancement from the anterior margin of the tumor leads to the radiologic characterization of this tumor as cystic renal carcinoma. We note that this enhancement is reproduced faithfully in the corticomedullary and nephrographic imputed images (Figure 4). Subject 5 displays a complex cystic tumor where the thickened margin is well seen in the imputed images (Figure 5).

**Figure 5.**
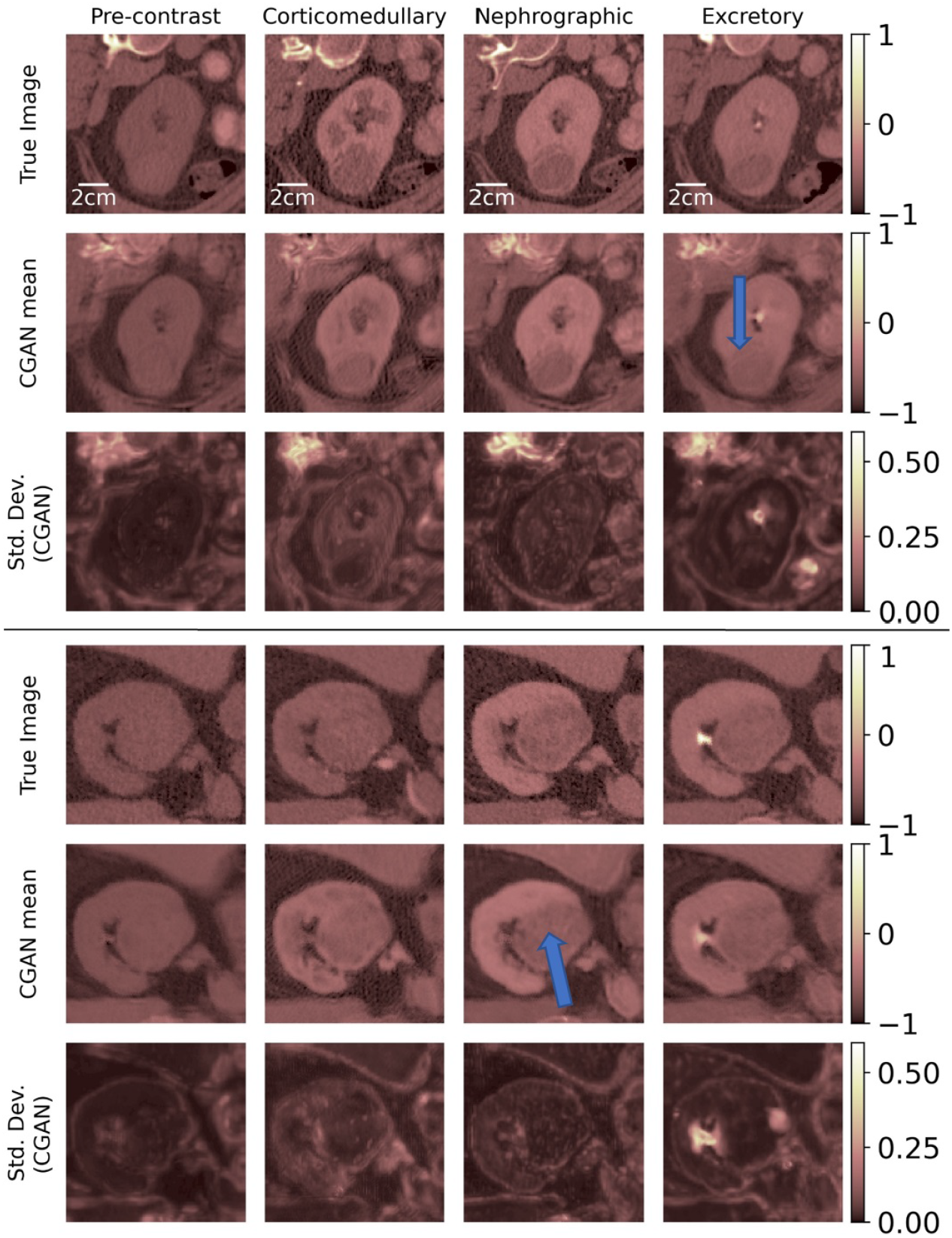
True and imputed imagesfor Subject 5 and 6.

Subject 6 displays a tumor that is a predominantly hypo-dense (low density compared to the adjacent normal renal tissue) with multiple internal components (arrow) which are well seen in the imputed images (Figure 5). This is a specific type of renal carcinoma, viz. papillary renal cell carcinoma where the density of the tumor is key to the diagnosis and is faithfully reproduced in the imputed images.

One of the distinguishing features of the method developed in this study is its ability to produce a distribution of imputed images conditioned on the images available at other time-points. This allows us to compute an image of the pixel-wise standard deviation for each imputed image. At the local level, the standard deviation image (shown in third row for each subject) highlights the regions where the uncertainty in the imputation is high. We note that the regions of high uncertainty tend to be at the interface between the kidney and abdominal cavity, and between other organs and the abdominal cavity. The total intensity of the standard deviation image also contains useful information.

In particular, as shown in Figure 6 we note that this value is good indicator of whether the NMSE of an imputed image is small or large. In particular, it can be used to classify (AUC = 0.88825) whether the NMSE for an imputed image will be above or below a threshold value of 0.1. This is particularly useful for downstream clinical tasks where imputed images with large standard deviation, and hence low-confidence, can be disregarded by the clinician.

**Figure 6.**
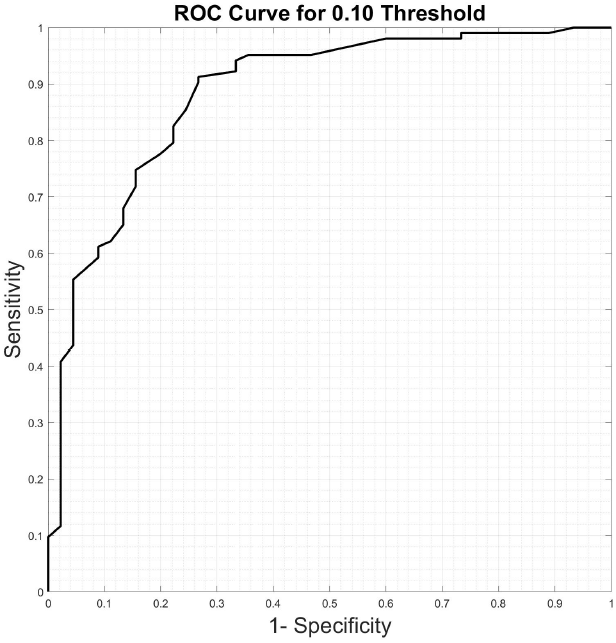
ROC curve for classifying a given imputed image as acceptable.

Finally, we note that even though the main advantage of our approach is to quantify the confidence in the imputation results, it also produces imaged with less error than other state of the art methods. In Table 1, its performance is compared with an enhanced PI2PIX approach that uses the same training and test data. For each time-point and in both metrics we observe that the cGAN based approach incurs smaller errors. Further in Table 2 it is compared with other state of the art methods, albeit on different training and test data. Here too, it quite clearly outperforms the other methods, even though these methods utilize additional information (like tumor segmented images) and contain losses that are much more complex. We believe that primary reason for the better performance of our approach is its stochastic nature. By virtue of this, the mean imputed image averages over any outlier predictions and does not let the outliers strongly influence the final estimate.

## Supporting Information

### Linear Regression

The linear regression prediction of each image is found by solving an over-determined system using the ordinary least-squares method. The predicted image is then the linear combination of the three other images that are available. Therefore, for example, for the pre-contrast phase, the linear regression prediction *R*^1^ is:

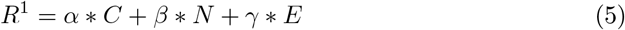

where C, N and E are the true corticomedullary, nephrographic and excretory images respectively. The coefficients *α, β, γ* are determined by minimizing the mean squared error between the predicted image and real image across all samples in the training set (without data augementation). The final linear maps obtained are expressed as

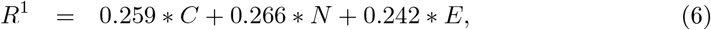

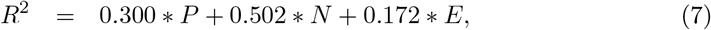

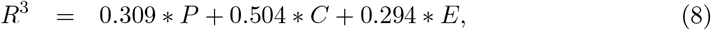

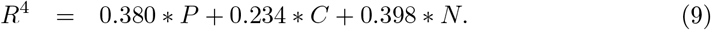

### Generator and Critic blocks

The generator and critic architectures are composed of several network blocks, as illustrated in Figure 2. The important components are explained below and while the training hyperparameters are listed in Table 3. In the generator we set *C* = 32, *k* = 16, *n* = 3, while in the critic we set *C* = 20, *k* = 16 and *n* = 2. The typical U-net uses Residual Blocks along with other neural networks. However, we have used Dense Blocks due to their superior performance. When it comes to the critic, a simple critic with downsampling has been used along with Dense Blocks. The standard layer normalization was used, unlike the generator where CIN was used. The exponential linear unit (ELU) was used in both the generator and the critic instead of the typical Leaky ReLU.

**Table 3.** Hyper-parameters for cWGAN.

| Hyper-Parameter | Value/Type |
| --- | --- |
| Training Samples | 12520 |
| Latent space dimension ($N_Z$) | 150 |
| Batch size | 15 |
| Number of training epochs | 800 |
| Number of $D$ updates per $G$ update | 4 |
| ELU activation parameter | 0.2 |
| Regularization parameter | 1e-7 |
| Optimization technique | Adam ($\beta_1 = 0.5$, $\beta_2 = 0.9$) |
| Learning rate | 0.001 |

### Convolution

The notation Conv(n, s, k) represents a 2D convolution operation in which k filters of size n are applied to the input tensor with a stride s. If n is greater than 1, reflective padding of width 1 is added to the input tensor in the spatial dimensions before the convolution. In case the third argument is not given, and the number of filters is set equal to the number of channels in the input tensor.

### Conditional Instance Normalization (CIN)

The idea behind CIN [8] is that the latent variable *z* is injected into the generator at all levels of the generator (U-net). The CIN block take as input the latent vector *z* and an intermediate tensor *w* of shape *H′ ×W×′ C′* and normalizes it along each channel *j* = 1, …, *C*^*′*^ in the following way:

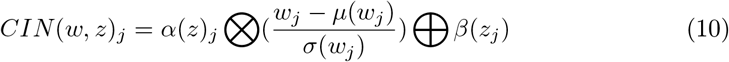

where 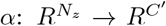 and 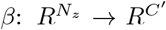 are learnable convolution layers that take *z* as input ⊗. and ⊕are used to represent mathematical operations. ⊗represents element-wise multiplication, which means that each element of one tensor is multiplied by the corresponding element of the other tensor ⊕ represents summation in the channel direction, which means that the values in each channel of the tensors are summed together. *µ*(*w*_*j*_) and *σ*(*w*_*j*_) compute the mean and fluctuations along the spatial directions, respectively. Therefore, each feature of the intermediate tensor is broken down into these mean and fluctuating components which are stochastic depending on the value of *z* through the learned layers *α* and *β*.

### Dense Block

DenseBlocks are a type of building block used in convolutional neural networks for image classification tasks. They were introduced in the DenseNet architecture, which is a deep neural network that connects each layer to every other layer in a feed-forward fashion. We use the notation DenseBlock(*k, n*), where *k* represents the number of filters of size 3 and stride 1 in the 2D convolution, and *n* represents the number of sub-blocks. An example of a DenseBlock with n equal to 3 is shown in Figure 7. When the DenseBlock appears in the generator, it receives two inputs: an intermediate tensor *w* and a latent variable *z*. The latent variable is incorporated into the block using conditional instance normalization (CIN). However, when the DenseBlock appears in the critic, it only receives the intermediate tensor w as input. In this case, CIN is replaced with layer normalization. In a DenseBlock, each layer receives the feature maps of all preceding layers as input, which are concatenated channel-wise. This means that the output of each layer in the DenseBlock is fed as input to all subsequent layers, allowing for highly efficient information flow throughout the network. By densely connecting the layers, DenseBlocks aim to reduce the vanishing gradient problem and improve gradient flow, which can lead to faster convergence during training.

**Figure 7.**
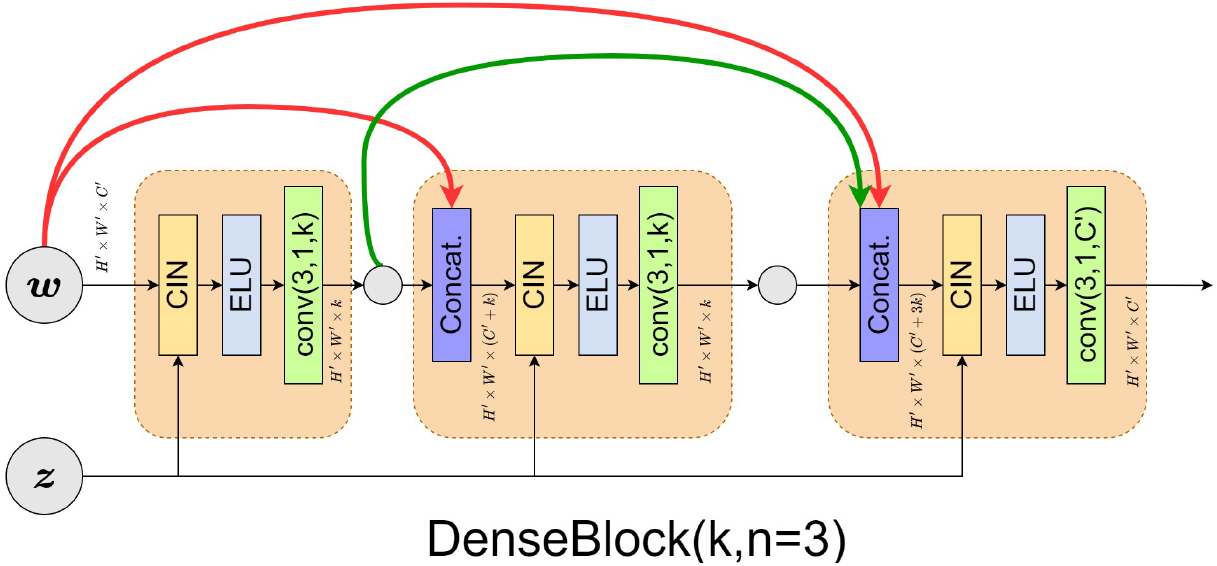
Architecture of a DenseBlock.

Overall, DenseBlocks are a powerful tool for building highly efficient and accurate convolutional neural networks, especially for image classification tasks with limited data. Dense Blocks therefore work on preserving the feed-forward nature of the network as each layer obtains additional inputs from all the layers preceding it and then in itself passes feature-maps to all subsequent layers.

### Down-Sampling Block

The down-sampling block (Figure 8) is designed to decrease the spatial resolution while simultaneously increasing the number of channels. This block comprises a convolution layer that produces an output with twice the number of channels as the input (*q* equal to 2), a 2D average pooling layer that decreases the spatial dimensions by a factor *p* of two, and a Dense block. The latent variable *z* is also an input when this block appears in the cGAN generator.

**Figure 8.**
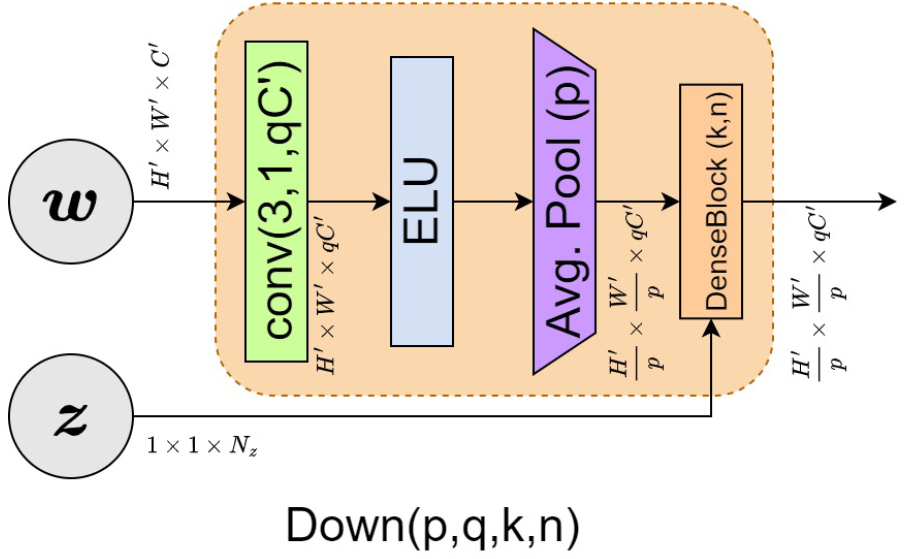
Architecture of a Down block.

### Up-Sampling Block

The up-sampling block (Figure 9) takes in the output **w** of the previous block along with the output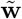 of a down-sampling block of the same spatial size via a skip connection.

**Figure 9.**
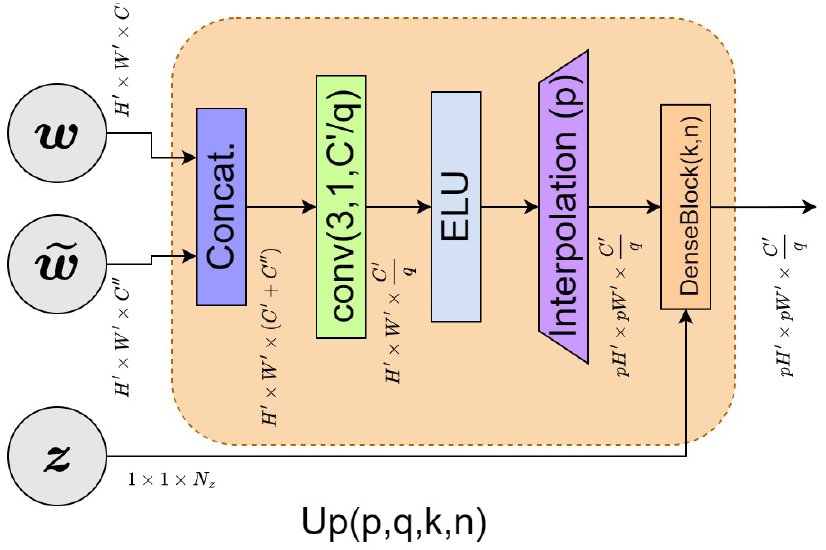
Architecture of an Up block.

These tensors are then concatenated together in the channel dimension. Next, the up-sampling block carries out a convolution operation that divides the channel size by *q* equal to 2, followed by a 2D up-sampling operation that doubles the spatial dimension. Here, a 2D nearest neighbour interpolation increases the spatial resolution by a factor *p* of 2. Finally, the signal is passed through a Dense block.

### The Gradient Penalty Term

The gradient penalty term is added to constrain the critic to be 1-Lipschitz on Ω_*X*_.

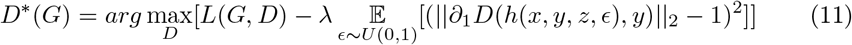

The derivative in the gradient penalty term denotes the derivative with respect to the first argument of the critic. Also, *U* (0, 1) represents the uniform distribution on [0, 1]. The function *h*(*x, y, z, ϵ*) is:

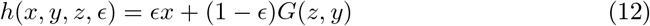

The gradient penalty parameter *λ* is set equal to 10.

## Acknowledgments

The support from ARO grant W911NF2010050 and the Ming-Hsieh Institute at USC is acknowledged.

